# Direct Virus-Bacteria Binding Enhances *Streptococcus equi* subsp. *zooepidemicus* Colonisation and Bacterial-Driven Immune Activation During H3N8 Equine Influenza A Virus Co-Infection

**DOI:** 10.64898/2026.05.19.724896

**Authors:** Askar K. Alshammari, Meshach Maina, Meshari A. Alsuwat, Adam M. Blanchard, Janet M. Daly, Stephen P. Dunham

## Abstract

Respiratory viral-bacterial co-infections cause severe disease across species, yet the molecular mechanisms underlying enhanced pathogenesis remain poorly understood. This study characterised H3N8 equine influenza A virus (IAV) and *Streptococcus equi* subspecies *zooepidemicus* (SEZ) co-infections using complementary ultrastructural and transcriptomic approaches. Transmission electron microscopy demonstrated direct physical binding between spherical (A/equine/Miami/63) and filamentous (A/equine/Sussex/89 and A/equine/Newmarket/5/2003) IAV isolates and SEZ, including when SEZ was heat-inactivated (θSEZ). Lectin staining revealed that SEZ expresses predominantly α2,3-linked sialic acids, the receptor for equine IAV. However, virus-bacteria binding persisted despite neuraminidase treatment. Scanning electron microscopy quantification demonstrated that viral pre-infection significantly enhanced bacterial adherence to cells of the DH82α canine macrophage-like cell line (2-fold increase, p<0.01) but not ExtEqFL (equine lung-derived) cells, revealing cell-type-specific enhancement. RNA-sequencing analysis showed that bacterial infection drove most transcriptional changes during co-infection with little difference in the number of differentially expressed genes (DEGs) between infection with SEZ alone (146 DEGS) or after pre-infection with either A/equine/Sussex/89 (166 DEGS) or A/equine/Newmarket/5/2003 (149 DEGS). Validation of upregulation of selected cytokines by RT-qPCR and ELISA demonstrated that SEZ infection drives dramatic cytokine upregulation compared to mock or θSEZ controls. Viral pre-infection did not alter the SEZ-induced pro-inflammatory cytokine responses (IL-6, IL-8, TNF-α) but significantly reduced IFN-β expression compared to SEZ infection alone. These findings suggest that direct virus-bacteria physical interactions may drive cell-type-specific enhancement of bacterial colonisation, fundamentally advancing our understanding of respiratory co-infection pathogenesis.

## 1. Introduction

Respiratory viral-bacterial co-infections represent a major clinical challenge across human and veterinary medicine, being consistently associated with increased disease severity, prolonged illness, and elevated mortality [1, 2]. The phenomenon of viral predisposition to secondary bacterial infections has been recognised for over a century, with the 1918 influenza pandemic serving as a stark historical reminder where bacterial pneumonia was the predominant cause of death [3]. Despite decades of research, the underlying mechanisms governing viral-bacterial synergy remain incompletely understood, particularly regarding direct molecular interactions between pathogens.

Current paradigms for the pathogenesis of co-infections focus primarily on indirect mechanisms, including virus-induced immunosuppression, epithelial barrier disruption, and impaired bacterial clearance [4, 5]. However, emerging evidence suggests that direct physical interactions between viral and bacterial pathogens may play crucial roles in enhanced pathogenesis. Studies in human respiratory infections have demonstrated direct binding between influenza A viruses and *Streptococcus pneumoniae*, mediated by shared recognition of sialic acid receptors [6-9]. These direct interactions can enhance bacterial adherence, increase virulence factor expression, and promote biofilm formation [9-11].

In equine medicine, respiratory co-infections involving equine influenza A virus (IAV) and bacterial pathogens such as *Streptococcus equi* subspecies *zooepidemicus* (SEZ) are frequently observed in clinical practice [12-14]. SEZ normally colonises the equine upper respiratory tract asymptomatically but can cause severe pneumonia and systemic complications, particularly following viral respiratory infections [13, 15]. The molecular mechanisms underlying this enhanced pathogenicity remain poorly characterised, representing a significant knowledge gap in understanding equine respiratory disease.

Equine IAV exhibits considerable genetic diversity, with Florida clade 2 isolates such as A/equine/Newmarket/5/2003 (N/5/03) possessing truncated non-structural 1 (NS1) and polymerase basic 1-F2 (PB1-F2) proteins compared to earlier isolates like A/equine/Sussex/1989 (Sussex/89) that encode full-length versions [16]. These differences significantly impact viral pathogenesis and immune evasion capabilities, with N/5/03 associated with more severe clinical outcomes despite reduced interferon antagonism [16, 17]. The biological relevance of these viral characteristics in cross-species contexts is well-documented, with multiple reports of H3N8 equine IAV transmission to canines, including racing greyhounds in Florida around 2004 [18] and English foxhounds in the United Kingdom in 2002 [19]. These interspecies transmission events resulted in severe respiratory disease with fatality rates of 5–8% in greyhounds, with fatal outcomes primarily attributed to co-infections with secondary bacterial pathogens, most notably SEZ strains cultured from affected animals [18]. The clinical significance of SEZ as an emerging pathogen in canine infectious respiratory disease is further supported by comprehensive reviews identifying it as a key bacterial contributor to complex respiratory syndrome in dogs [20]. This documented association between equine-origin H3N8 influenza virus and SEZ co-infection in canines, combined with demonstrated receptor compatibility of canine respiratory tissue for equine influenza virus infection [19], provides compelling biological justification for using canine DH82α cells to investigate IAV-SEZ co-infection dynamics.

Host transcriptional responses during viral-bacterial co-infections differ markedly from single-pathogen infections, with evidence suggesting complex interplay between viral immune evasion mechanisms and bacterial inflammatory drivers [21-23]. Macrophages serve as critical first responders to respiratory pathogens and play pivotal roles in co-ordinating innate immune responses [24, 25]. Understanding the cell-type-specific responses to co-infection is essential for elucidating pathogenic mechanisms and identifying therapeutic targets.

Our comprehensive scoping review of molecular interactions between influenza A virus and Streptococcus proteins in co-infection contexts revealed significant knowledge gaps regarding direct pathogen interactions and their functional consequences [26]. Most studies have focused on temporal sequences of infection rather than simultaneous pathogen interactions, highlighting the need for integrated approaches combining structural and functional analyses.

This study aimed to characterise the molecular and cellular mechanisms underlying equine IAV-SEZ co-infection using complementary ultrastructural and transcriptomic approaches. We employed transmission electron microscopy (TEM) to visualise direct virus-bacteria interactions, scanning electron microscopy (SEM) to quantify enhanced bacterial adherence, and RNA sequencing (RNA-seq) to define host transcriptional responses during co-infection. Our findings highlight potential mechanisms of viral–bacterial synergy that may contribute to respiratory co-infection pathogenesis.

## 2. Materials and methods

### 2.1 Cell lines and influenza A viruses

Canine macrophage-like DH82α cells were obtained from ECACC (UK), and ExtEqFL (equine lung) cells were kindly provided by Prof. Pablo Murcia (University of Glasgow, UK). Both cell types were maintained in Dulbecco’s Modified Eagle Medium (DMEM) (Gibco, UK), supplemented with 10% fetal calf serum, 100 U/ml penicillin, and 100 μg/ml streptomycin at 37°C in 5% CO_2_.

Three equine H3N8 IAV isolates were used: A/equine/Miami/1963 (Miami/63), A/equine/Sussex/89 (Sussex/89), and A/equine/Newmarket/5/2003 (N/5/03). All viruses were propagated in 10-day-old embryonated hens’ eggs, and allantoic fluid was harvested at 72 h post-inoculation. Viruses were titrated using the 50% tissue culture infectious dose (TCID_50_) method in DH82α cells based on Spearman-Kärber [27]. Viral stocks were aliquoted and stored at -80°C until further use.

### 2.2 Bacterial strain and growth conditions

The D2a strain of *Streptococcus equi* subsp. *zooepidemicus* (SEZ), originally isolated from a horse with pneumonia, was obtained from Dr Andrew Waller (Animal Health Trust, UK). *Streptococcus suis* (*S. suis*) P1/7, a reference type strain (GenBank accession number AM946016.1) originally isolated from a case of clinical meningitis in a pig, represents a typical serotype 2 strain, the most common serotype identified in both human and porcine infections [28]. Both SEZ and *S. suis* were cultured on Todd-Hewitt broth (THB) supplemented with 10% fetal calf serum at 37°C. Bacterial concentrations were determined by colony-forming unit per ml (CFU/ml) [29].

### 2.3 Heat-inactivated bacteria

SEZ and *S. suis* were heat-inactivated by incubation in a water bath at 70°C for 30 min. Following heat treatment, bacterial suspensions were allowed to cool to room temperature before further use. Complete inactivation was determined by plating aliquots of the treated suspensions onto Todd-Hewitt Agar (THA) and incubating overnight at 37°C, to confirm the absence of bacterial colonies.

### 2.4 Transmission electron microscopy

To assess virus-bacteria binding, 1×108 CFU/ml of bacteria were harvested by centrifugation (5,000 × g, 10 min) and resuspended in 1 ml of phosphate buffered saline (PBS) containing 1×10^5^ TCID_50_ virus. The mixtures were incubated at room temperature for 30 min, 1 h, or 2 h to determine the optimal incubation period. Based on preliminary observations, 1 h incubation produced the most distinct viral attachment and optimal structural preservation and was therefore used for all subsequent experiments.

Following incubation, samples were washed three times with PBS by centrifugation at 10,000 × g for 3 min to remove unbound viral particles. The pellet was resuspended in 1 ml of PBS. Then, 10 µl of the suspension was applied to continuous carbon-coated 300-mesh copper grids (EM Resolution, UK) and allowed to adsorb for 10 min. Excess liquid was removed using filter paper. The grids were then exposed to ultraviolet (UV) light (254 nm, 20 min; Flowgen, UK) to ensure complete viral and bacterial inactivation [30]. Negative staining was performed by applying 10 µl of 2% phosphotungstic acid (PTA) at pH 7.4 (Sigma, UK) to the grids for 30 s. Excess stain was removed using filter paper, and grids were air-dried at room temperature for 30 min. Images were further analysed using Amira software (Thermo Scientific, UK) to highlight viral structural components and genome.

### 2.5 Lectin staining

Lectin staining of bacterial surfaces was performed to characterize sialic acid linkage patterns. Briefly, 1×10^7^ CFU of SEZ and *S. suis* were applied to Epredia Superfrost Plus adhesion microscope slides (Thermo Fisher Scientific, UK) and air-dried at room temperature. Slides were treated with acetone for 10 min at 20°C, washed twice with 1 x Tris-buffered saline (TBS), and blocked sequentially with streptavidin and biotin blocking solutions (Vector Laboratories, UK) for 15 min.

Two biotinylated lectins were used to detect specific sialic acid linkages: *Maackia amurensis* lectin II (MAL-II) for α2,3-linked sialic acids and *Sambucus nigra* lectin (SNA) for α2,6-linked sialic acids (Vector Laboratories, UK). Both lectins were diluted 1:200 in TBS and incubated overnight at 4°C in a dark humid chamber. Following three 5-min TBS washes, slides were incubated with streptavidin-Alexa-Fluor594 (1:500 dilution; Thermo Fisher, UK) for 2 h. After final TBS washes, samples were mounted using ProLong Gold with DAPI (Invitrogen, UK) and cured for 24 h in darkness at room temperature. Samples were observed and imaged using fluorescence microscopy (Leica).

### 2.6 Neuraminidase treatment of bacteria

Live and heat-inactivated (θSEZ) and *S. suis* were harvested by centrifugation at 5,000 × g for 10 min and washed twice with PBS. The bacterial pellets were resuspended in PBS to a final concentration of approximately 1×10^6^ CFU per 50 μl. Bacteria were treated with either neuraminidase type V from *Clostridium perfringens* (Sigma-Aldrich, UK) or neuraminidase (sialidase) from *Vibrio cholerae* (Roche, Switzerland) at a final activity of 400 mU per 50 µl reaction volume. The mixtures were incubated at 37°C for 1 h with gentle mixing. Following enzymatic digestion, reaction mixtures were centrifuged at 5,000 × g for 10 min and supernatants discarded. Bacterial pellets were washed three times with cold PBS, centrifuging at 5,000 × g for 10 min between washes to remove residual enzyme. Neuraminidase-treated bacteria were resuspended in PBS and used immediately for subsequent binding assays. Untreated bacteria processed in parallel without neuraminidase served as controls. This protocol was adapted from Nita-Lazar et al. [31] with modifications.

### 2.7 Scanning electron microscopy and quantitative analysis

DH82α and ExtEqFL cells were seeded at 5×10^5^ cells/well on borosilicate glass coverslips (Ø 19 mm; VWR, UK) in 12-well plates and grown overnight in DMEM supplemented with 10% FCS. Cells were infected in three biological replicates with one of three equine IAV isolates (Miami/63, Sussex/89, or N/5/03) at 1×104 TCID_50_/ml in serum-free DMEM containing 1 µg/ml TPCK-treated trypsin. After 24 h, cells were washed twice with PBS and subsequently infected with SEZ at 1×10^7^ CFU/mL for 1 h at 37°C.

Control groups included SEZ alone (without prior viral infection), uninfected cells, and virus-only controls.

Following infection, cells were fixed with 3% glutaraldehyde in 0.1 M cacodylate buffer at 4°C for 24 h, then washed twice with cacodylate buffer. Samples were dehydrated through a graded ethanol series (30%, 50%, 70%, 90%, and 100%) with 10-min incubations at each step, followed by overnight drying with hexamethyldisilazane (HMDS; Sigma-Aldrich, UK). Dried coverslips containing the fixed and dehydrated cells were carefully placed onto the double-sided carbon tape, ensuring full contact between the sample surface and the stub. Gentle pressure was applied to secure the coverslip without damaging the sample. Following mounting, a thin layer of platinum was applied using a sputter coater (agar auto sputter coater, Agar Scientific Ltd, UK) to enhance surface conductivity and minimise charging during imaging. The sputter coating was performed under vacuum conditions with argon gas introduced at a pressure of 0.3 mbar. Argon ions bombarded the carbon target, dislodging carbon atoms that were deposited evenly onto the sample surface. A coating thickness of approximately 14 nm was applied over a sputtering duration of 50 s.

Samples were examined using a JEOL IT-200 scanning electron microscope at 5.0 kV accelerating voltage, 11.5 mm working distance, and ×6000 magnification. Bacterial adherence was quantified using ImageJ software (version 1.54m, National Institutes of Health, USA)[32] by counting SEZ adhering to host cells across 60 randomly selected fields per condition (n=3 biological replicates). Statistical analysis was performed as described in section 2.12.

### 2.8 IAV and SEZ Co-infection of DH82α cells

Twelve-well plates were seeded with DH82α cells at a density of 5 × 10^5^ cells per well in 1 ml of DMEM supplemented with 10% FCS, in three biological replicates, and incubated overnight at 37°C with 5% CO_2_ until they reached approximately 80–90% confluency. The experimental design for viral, bacterial, and co-infection studies is illustrated in Figure 1.

**Figure 1.**
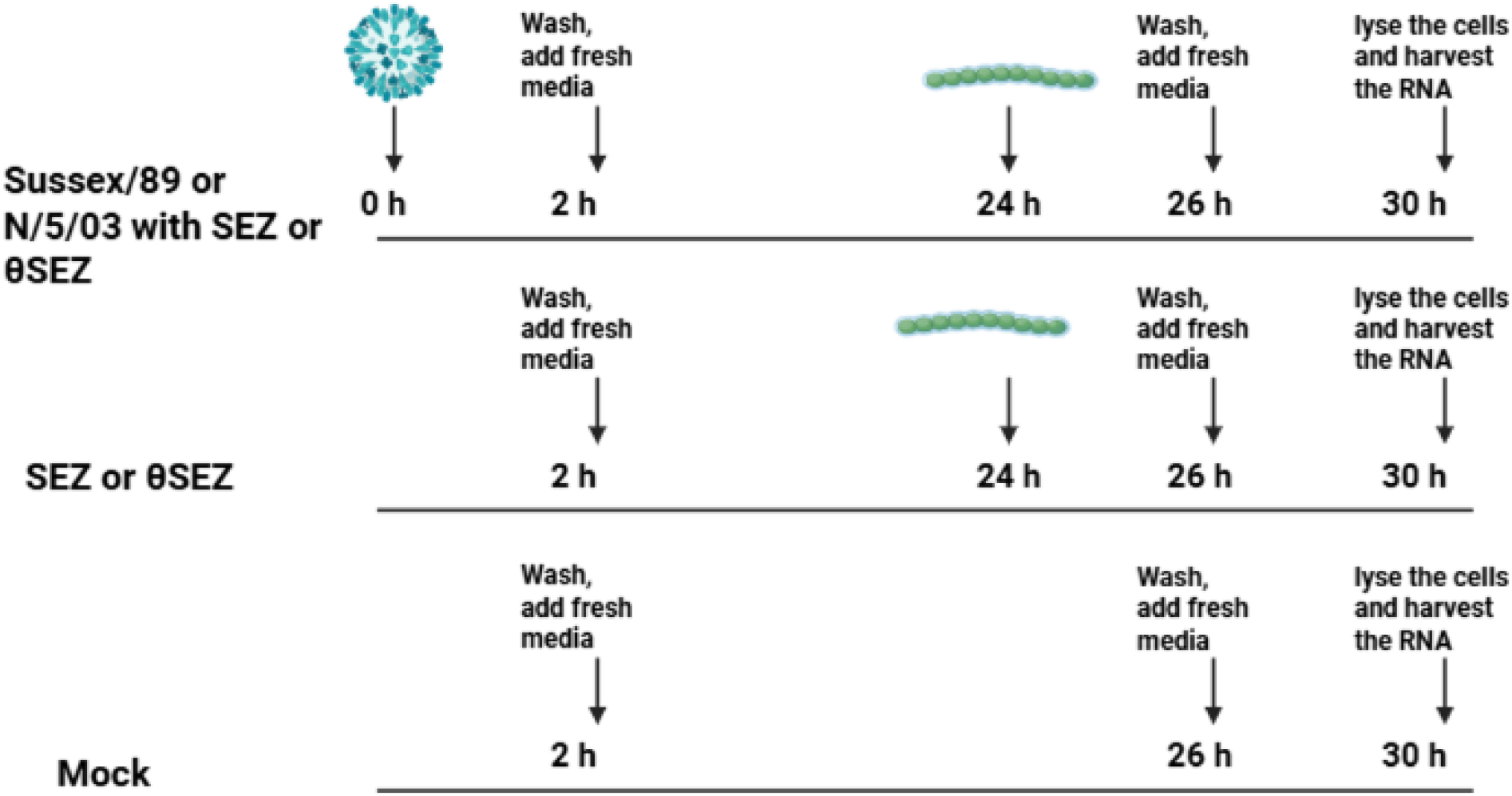
Experimental timeline for equine influenza A and *Streptococcus equi zooepidemicus* (SEZ) co-infection studies in DH82α cells. Equine influenza isolates: A/equine/Sussex/89 (Sussex/89), and A/equine/Newmarket/5/2003 (N/5/03). θSEZ: heat-inactivated SEZ.

For co-infection studies, cells were first inoculated with 1×104 TCID_50_/ml of Sussex/89 or N/5/03 in serum-free infection medium containing 1 μg/ml TPCK-trypsin. Following 2 h incubation, the inoculum was removed, cells were washed three times with PBS, and infected cells were incubated for 24 h at 37°C with 5% CO_2_ prior to bacterial infection.

Cells were infected with either SEZ or θSEZ at 1 × 10^5^ CFU/ml for 2 h, washed three times with PBS, and incubated for a further 6 h at 37°C with 5% CO_2_. Co-infections with SEZ were performed using both Sussex/89 or N/5/03, whereas co-infections with θSEZ were limited to N/5/03. Parallel infections were conducted with SEZ or θSEZ alone (without prior viral infection) under identical conditions, while mock-infected controls were incubated in infection medium only. Cells from all treatment groups were lysed at 30 h post-infection for RNA extraction and RNA-sequencing.

### 2.9 RNA sequencing and bioinformatics analysis

Total RNA was extracted from co-infected and control DH82α cells using a RNeasy Kit (Qiagen, UK) following the manufacturers protocol. RNA concentration was measured using a NanoDrop 8000 spectrophotometer, and quality was assessed using a Qubit 4 fluorometer (Thermo Fisher Scientific). Only samples with RNA Integrity Number (RIN) ≥ 7 were used for sequencing.

Library preparation and long non-coding RNA (lncRNA) sequencing were performed by Novogene. Briefly, ribosomal RNA was depleted from total RNA (≥ 500 ng), followed by ethanol precipitation and mRNA fragmentation. Directional cDNA libraries were constructed through first-strand synthesis with random hexamer primers, second-strand synthesis incorporating dUTPs, end repair, A-tailing, adapter ligation, size selection, and amplification. Sequencing was performed on an Illumina NovaSeq 6000 platform, generating 150 bp paired end reads, with approximately 9GB raw data per samples. Raw sequencing data are deposited in the NCBI Sequence Read Archive under BioProject accession PRJNA1454118 (SRR38177454–SRR38177459).

Quality assessment was performed using FastQC [33], with comprehensive monitoring via MultiQC (version 1.22.3) [34]. Read preprocessing was conducted using FastP (version 0.23.4) [35] to remove adapters and filter low-quality reads. Processed reads were aligned to the Labrador retriever transcriptome reference (ROS_Cfam_1.0, GCA014441545.1) using Salmon (version 1.10.1) [36].

Differential gene expression analysis was performed in R (version 4.4.1) using DESeq2 (version 1.44.0) [37] with significance thresholds set at adjusted p < 0.05, > log 2-fold change. Data visualisation included principal component analysis, heatmaps (pheatmap package), and volcano plots (EnhancedVolcano package). Pathway enrichment analysis was conducted using Gene Ontology (GOstats package) and KEGG databases (clusterProfiler package version 4.12.0). All analysis scripts are publicly available at: https://github.com/Askaralshammari/EIV-and-SEZ-RNAseq.

### 2.10 Quantitative RT-PCR

Selected genes identified by RNA-seq analysis were validated by quantitative RT-PCR. Total RNA (0.5–1 µg) was reverse-transcribed into complementary DNA (cDNA) using the SuperScript™ IV Reverse Transcriptase kit (Fisher Scientific, UK). Expression of host genes was quantified by RT-qPCR using a CFX Connect Real-Time PCR Detection System (Bio-Rad) with the SensiFAST SYBR No-ROX Kit (Bioline, UK). Gene-specific primers were used to measure the expression of selected cytokines and immune related genes: IL-6, TNF-α, IL-8, TLR2, TLR3, IFN-α, and IFN-β (Table 1). Hydroxymethylbilane synthase (HMBS) and 18S ribosomal RNA were used as reference genes. All the primers were designed for this study, except those for IFN-α [38], IFN-β [39], 18S, and HMBS [40].

**Table 1.**
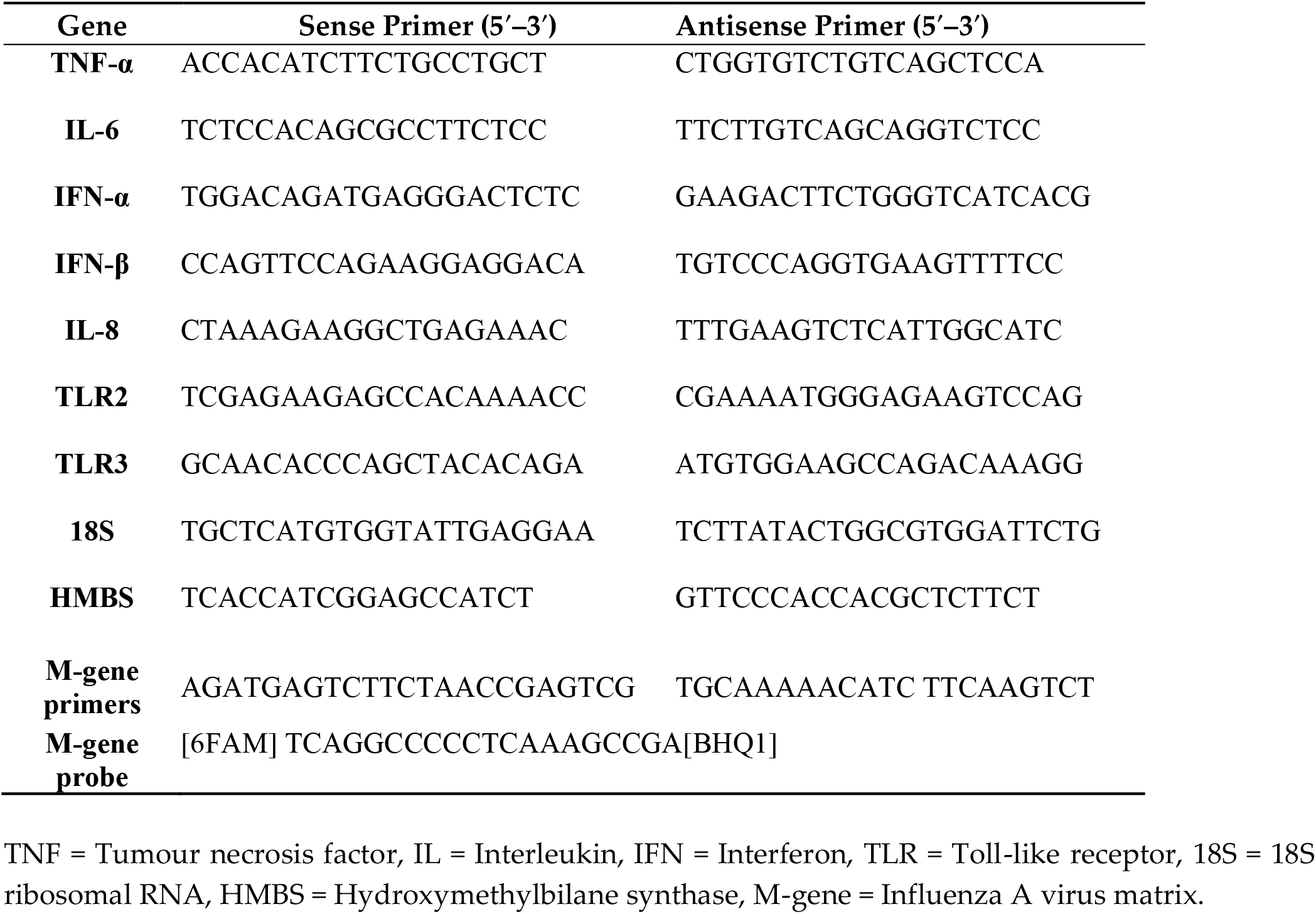
Primers used for RT-qPCR amplification.

Cycling conditions included initial polymerase activation at 95°C for 2 min, followed by 40 cycles of denaturation at 95°C for 5 s, annealing at 60°C for 10 s, and extension at 72°C for 10 seconds. Thereafter, melt curve analysis was performed from 65°C to 95°C with data collection every 2.2°C. Standard curves were generated using serial dilutions of pooled cDNA samples to calculate amplification efficiency for each primer set. Relative expression levels were calculated using the ΔΔCt method [41] with normalisation to the geometric mean of the two reference genes.

Viral RNA was extracted from DH82α cells using an RNeasy Kit (Qiagen, UK) following the manufacturer’s instructions. Influenza A virus M gene copy numbers were quantified using one-step RT-qPCR with the SensiFAST Probe No-ROX Kit (Bioline, UK). Primers and probe sequences targeting a 101 bp region of the M gene were used as described by Spackman et al. [42] and are detailed in Table 1. Each 20 μl reaction contained 16 μl of reaction mix and 4 μl of RNA template in 96-well plates (Thermo Scientific). Cycling conditions comprised: reverse transcription at 50°C for 30 min, initial polymerase activation at 95°C for 2 min, followed by 40 cycles of denaturation at 95°C for 15 s and annealing/extension at 60°C for 1 min. A standard curve was generated using serial dilutions of *in vitro*-transcribed M gene viral RNA to calculate absolute viral RNA levels. All samples were analysed in triplicate, and results were expressed as log_10_ M gene copies per μl.

### 2.11 Enzyme-Linked Immunosorbent Assays

Cytokine protein concentrations in cell culture supernatants were measured using commercial ELISA kits according to manufacturer protocols. Canine IL-6 and TNF-α were quantified using R&D Systems kits, while canine IFN-β was measured using RayBio kits (Cambridge Bioscience, UK). Absorbance was measured at 450 nm using a GloMax® Explorer microplate reader (Promega, UK), and cytokine concentrations were calculated from standard curves generated for each assay.

### 2.12 Statistical analysis

All experiments were performed in biological triplicate unless otherwise specified. Statistical analyses were conducted using GraphPad Prism version 10.0. Statistical significance was determined using appropriate tests as indicated in figure legends, with p<0.05 considered significant. Data are presented as mean ± standard deviation or standard error of the mean as indicated in figure legends.

For RNA-seq analysis, transcripts with a false detection rate corrected adjusted p < 0.05, > log 2-fold change greater than 1 and less than minus 1 were considered to be differentially expressed, unless otherwise stated. False detection rate correction was performed using the Benjamini-Hochberg method [43].

For SEM bacterial adherence quantification, a one-way ANOVA with Dunnett’s multiple comparisons test was used to compare treatment groups to untreated SEZ.

## 3. Results

### 3.1 Direct physical interactions between equine influenza A viruses and SEZ

TEM showed that the morphology of the virus isolates was consistent with previous morphological studies [44]: Miami/63 exhibited predominantly spherical virions (Figure 2A), while both Sussex/89 and N/5/03 displayed filamentous morphology (Figure 2B).

**Figure 2.**
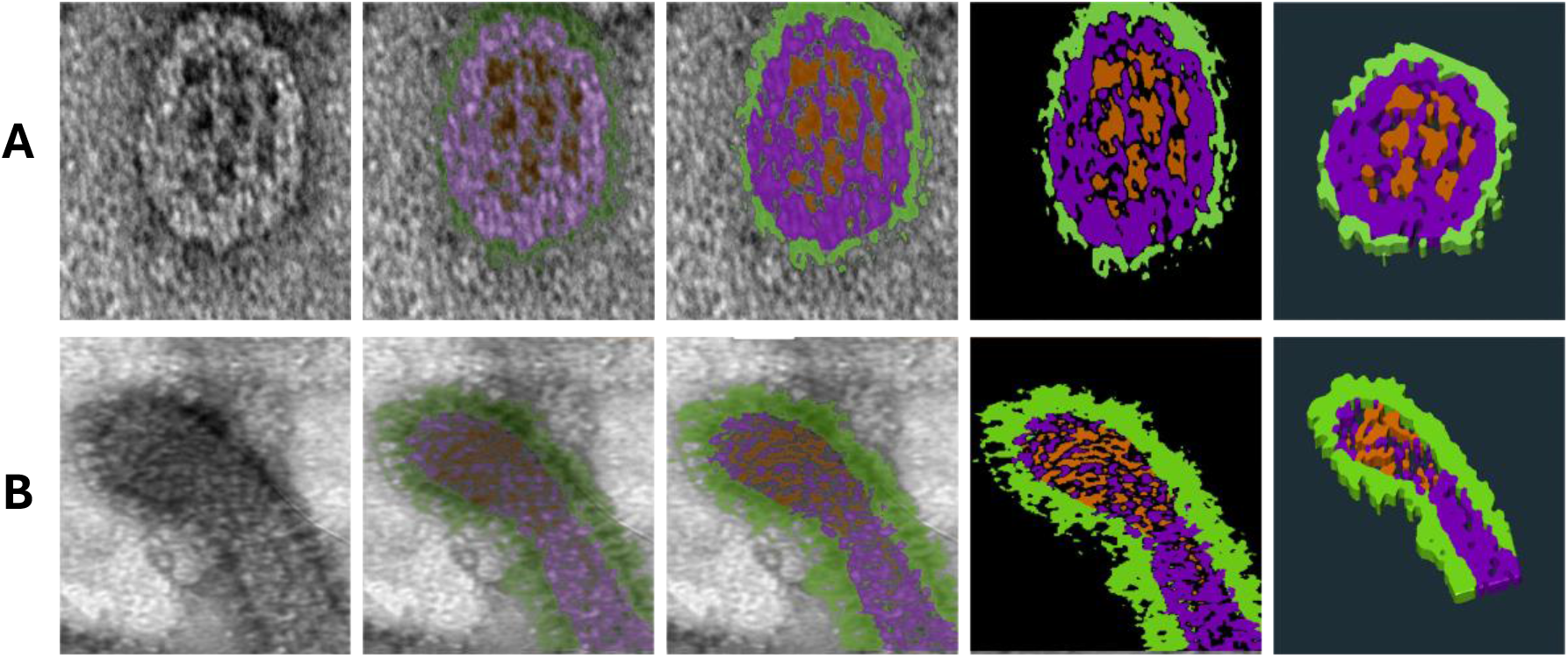
Transmission electron microscopy images showing the morphology of equine influenza virions. (A) Spherical virion representing Miami/63. (B) Longitudinal section of the tip of a filamentous virion representative of Sussex/89 and N/5/03. Images were manually segmented and colour-enhanced using Amira software (Thermo Scientific, UK) to highlight structural components: viral glycoproteins (green), membrane and associated matrix (purple), and viral genome (brown).

TEM revealed direct physical interactions between equine IAV particles and both SEZ and *S. suis* (Figure 3). All three isolates examined demonstrated clear attachment to bacterial surfaces, regardless of bacterial activity. Viral binding was consistently observed with both active and heat-inactivated bacteria, indicating that the interaction does not require active bacterial surface proteins or other structures susceptible to heat denaturation (Figure 3).

**Figure 3.**
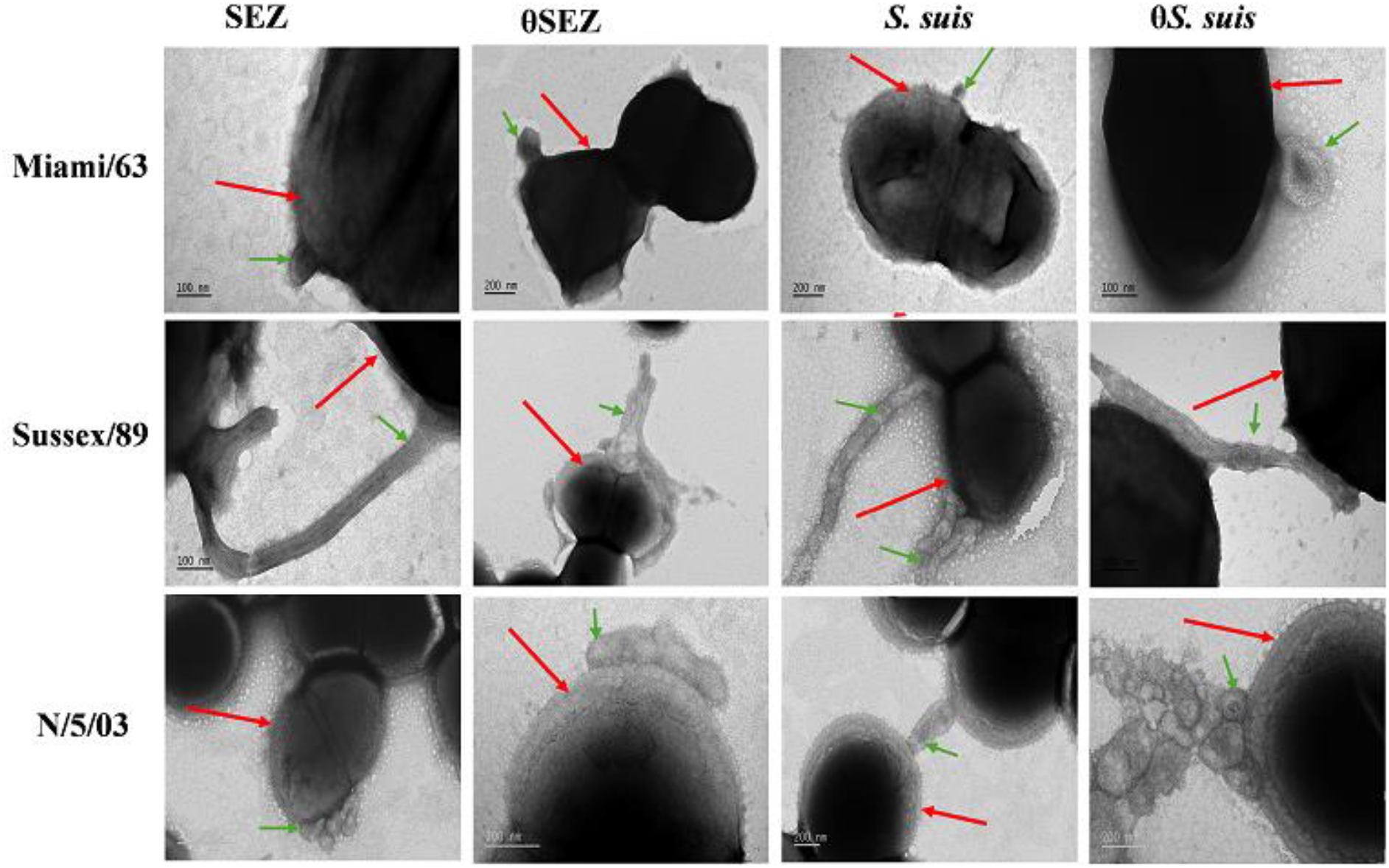
Direct interaction between equine influenza A viruses and streptococcal bacteria. Transmission electron microscopy images showing equine influenza A virus isolates Miami/63, Sussex/89 or N/5/03 bound to active or heat-inactivated (θ) *Streptococcus equi* subspecies *zooepidemicus* (SEZ) and *Streptococcus suis (S. suis)*. Green arrows: virus particles; red arrows: bacterial surfaces. Scale bars: 100–200 nm.

Morphological differences between virus isolates influenced the nature of physical interactions. Filamentous virus isolates (Sussex/89 and N/5/03) exhibited extensive contact with bacterial surfaces along their length, while the predominantly spherical Miami/63 isolate showed point-contact attachment to bacteria. Notably, viral particles were frequently observed in close proximity to bacterial cells, with individual virions appearing to bridge between multiple bacteria in some instances.

Neuraminidase treatment of the bacteria did not abrogate Sussex/89 binding to either SEZ or *S. suis* surfaces, regardless of bacterial activity (data not shown).

### 3.2 Surface sialic acid expression differs between SEZ and *S. suis*

To investigate the potential role of sialic acids in virus-bacteria interactions, lectin staining was performed to characterise the sialic acid linkage patterns surfaces of SEZ and *S. suis* bacteria. The two bacterial species exhibited markedly different sialic acid expression profiles (Figure 4). SEZ displayed predominantly α2,3-linked sialic acids, as evidenced by strong positive staining with *Maackia amurensis* lectin II (MAL-II; red fluorescence), while showing minimal binding of *Sambucus nigra* agglutinin (SNA; green fluorescence). In contrast, *S. suis* showed abundant α2,6-linked sialic acids detected by strong SNA staining, with negligible MAL-II binding. Neuraminidase treatment to remove bacterial surface SA did not affect viral binding to either bacterial species; Sussex/89 virions remained attached to both active and inactive bacteria following neuraminidase treatment (data not shown).

**Figure 4.**
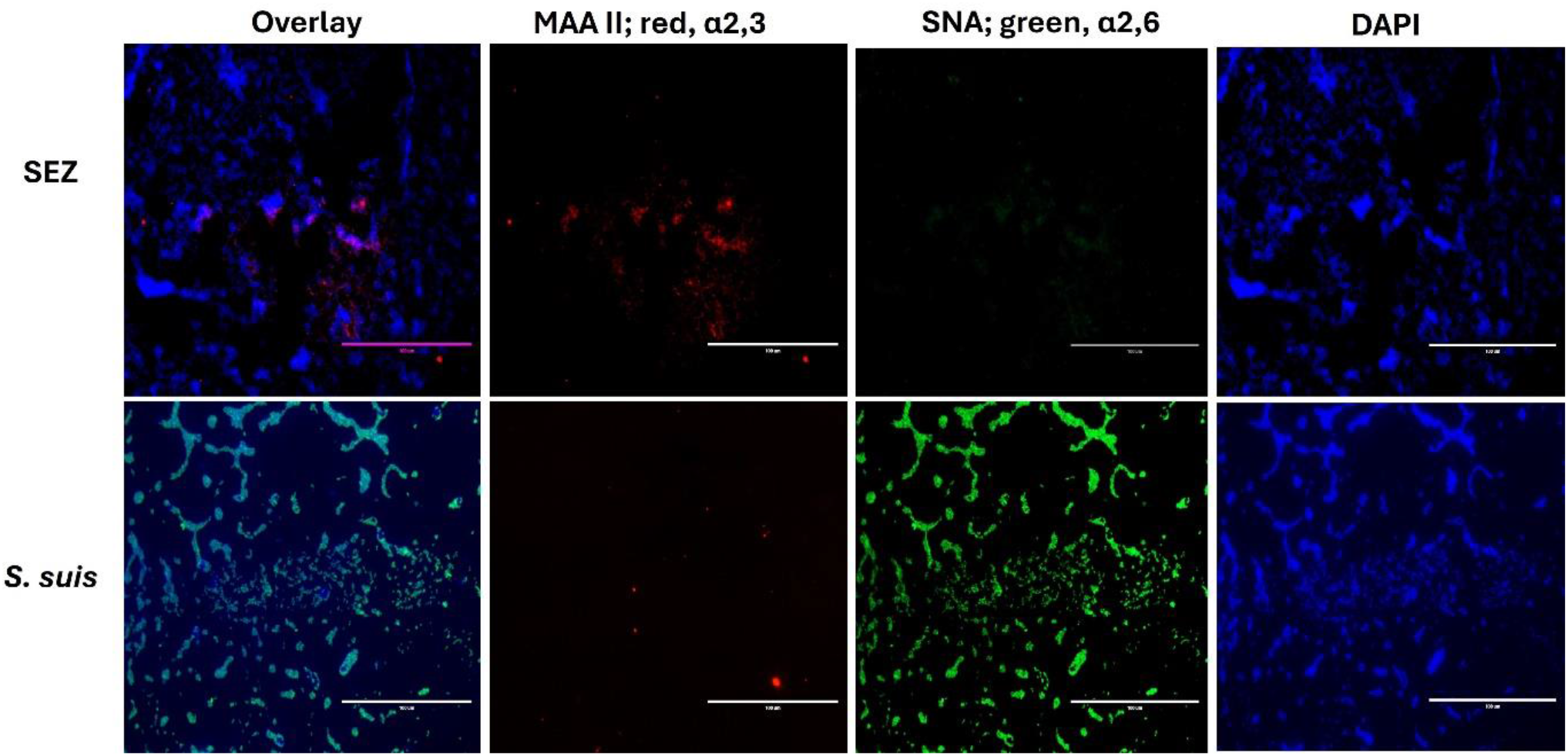
Differential sialic acid linkage patterns on *Streptococcus equi* subspecies *zooepidemicus* (SEZ) and *Streptococcus suis (S. suis)* bacterial surfaces. Fluorescence microscopy of lectin staining showing sialic acid linkage specificity on bacterial surfaces. Top row: SEZ; Bottom row: *S. suis*. Bacteria were stained with *Maackia amurensis* lectin II (MAL-II; red) specific for α2,3-linked sialic acids and *Sambucus nigra* agglutinin (SNA; green) specific for α2,6-linked sialic acids. DAPI (blue) shows bacterial nuclei. Overlay shows merged fluorescence channels. Scale bars 100 μm.

### 3.3 Cell-type specific enhancement of bacterial adherence following viral pre-infection

SEM imaging revealed that SEZ bacteria adhered to the surface of both DH82α and ExtEqFL cells across all experimental conditions, appearing as individual cocci or in characteristic streptococcal chain formations (Figure 5A). Adherent bacteria were clearly visible on cell surfaces, with bacterial chains frequently observed extending across the cell surface. Mock-infected cells displayed normal cell surface morphology without adherent bacteria, confirming the specificity of bacterial attachment.

**Figure 5.**
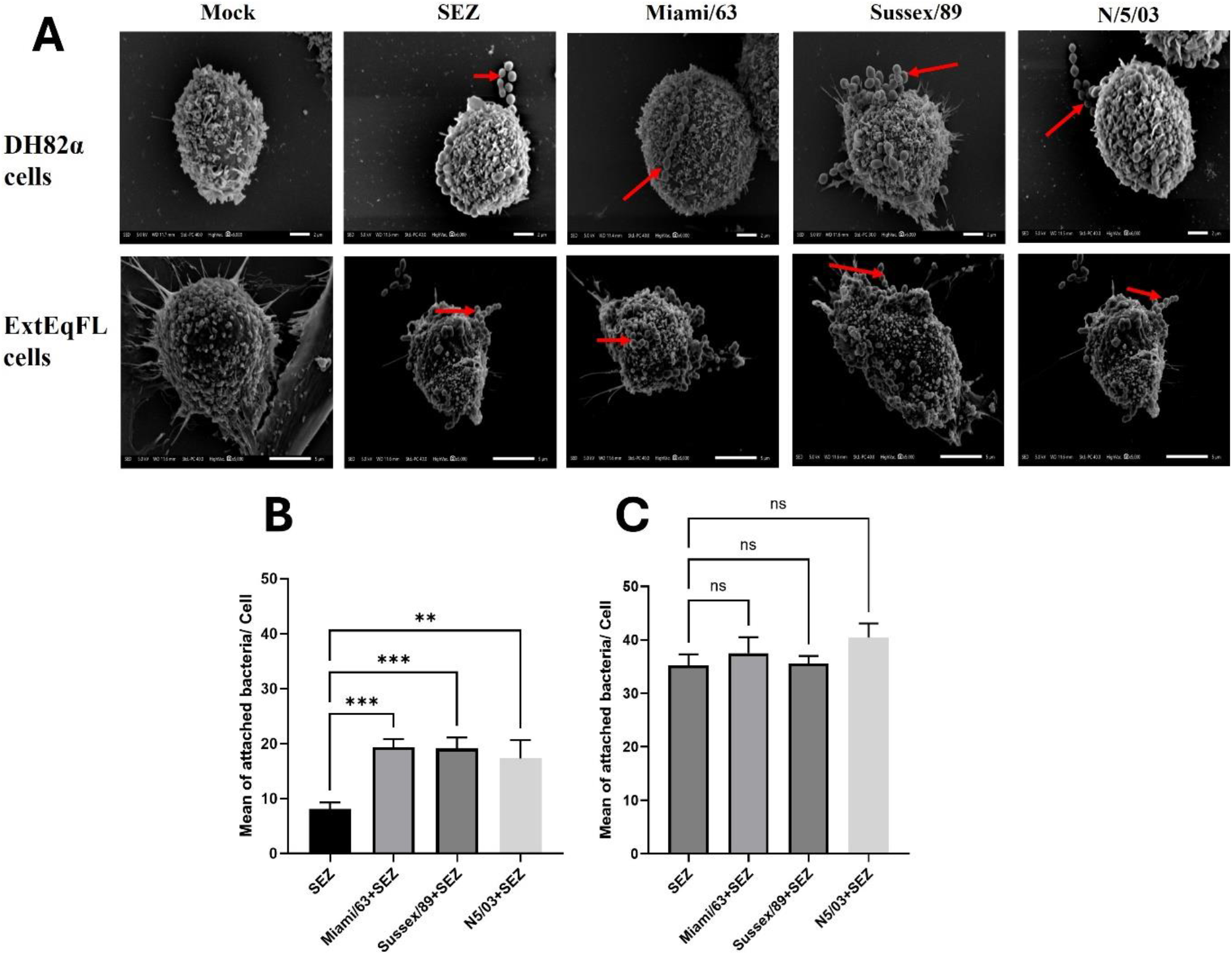
Scanning electron microscopy (SEM) reveals cell-type-specific enhancement of *Streptococcus equi* subspecies *zooepidemicus* (SEZ) adherence following viral pre-infection. (A) Representative SEM images of DH82α cells (top row) and ExtEqFL cells (bottom row) pre-infected with equine IAV isolates (Miami/63, Sussex/89, N/5/03) for 24 h, followed by incubation with SEZ for 1 h. Red arrows indicate adherent SEZ bacteria on cell surfaces, appearing as individual cocci or characteristic streptococcal chains. Mock-infected cells display normal cell morphology without bacterial exposure. SEZ without virus shows bacterial attachment to cells without prior viral infection. Scale bars = 2 μm (DH82α cells) and 5 μm (ExtEqFL cells). (B) Quantification of adherent SEZ bacteria per DH82α cell. Viral pre-infection with all three isolates significantly increased bacterial attachment compared to SEZ alone (approximately 2-fold increase). (C) Quantification of adherent SEZ bacteria per ExtEqFL. No significant differences were observed between viral pre-infection conditions and SEZ alone control. Bacterial counts were determined from SEM images by manual counting using ImageJ software (60 fields of view per replicate, three biological replicates per condition). Data show mean ± SD. **p < 0.01; ***p < 0.001 vs. SEZ alone; ns, not significant (one-way ANOVA with Dunnett’s multiple comparisons test).

Quantification of adherent bacteria from SEM images demonstrated cell-type-specific responses to viral pre-infection. In DH82α cells, all three virus isolates significantly increased bacterial attachment compared to SEZ alone (Figure 5B). This represents an approximately 2-fold increase in bacterial adherence following viral infection, with no significant differences observed between the three virus isolates. In contrast, although a greater number of bacterial cells were observed to adhere to ExtEqFL cells, likely reflecting their larger size, this was not significantly enhanced by viral pre-infection (Figure 5C).

These findings demonstrate that prior infection with IAV can significantly enhance bacterial adherence in a cell-type-dependent manner, suggesting that host cell characteristics influence the functional consequences of virus-bacteria interactions during co-infection.

### 3.4 Differential host gene expression during EIV-SEZ co-infection

RNA-seq analysis was performed on DH82α cells under six experimental conditions: (i) mock infection, (ii) SEZ alone, (iii) θSEZ alone, and co-infections of (iv) Sussex/89 and SEZ, (v) N/5/03 and SEZ or (vi) N/5/03 and θSEZ. Principal component analysis revealed that bacterial activity was the primary determinant of transcriptional responses (Figure 6A), with SEZ alone and both co-infections (Sussex/89 and SEZ and N/5/03 and SEZ), clustering closely together, indicating similar global gene expression profiles. In contrast, θSEZ alone, or with N/5/03, and mock samples formed a separate cluster, demonstrating that bacterial activity substantially influences the host transcriptional response during co-infection. Differential gene expression analysis confirmed this pattern (Figure 6B), with SEZ alone inducing 146 differentially expressed genes (99 upregulated, 47 downregulated) compared to mock controls. The numbers of DEGs were similar for co-infection with Sussex/89 (166 DEGs: 104 upregulated, 62 downregulated) or N/5/03 (149 DEGs: 93 upregulated, 56 downregulated). The θSEZ produced no significant transcriptional changes and N/5/03 co-infected with θSEZ yielded only 9 DEGs (3 upregulated, 6 downregulated). Venn diagram analysis revealed substantial overlap, with 76 genes (59.4%) commonly upregulated across SEZ alone or with Sussex/89 or N/5/03 pre-infection (Figure 6C). Analysis of downregulated genes showed less overlap, with 29 genes (34.1%) commonly downregulated (Figure 6D).

**Figure 6.**
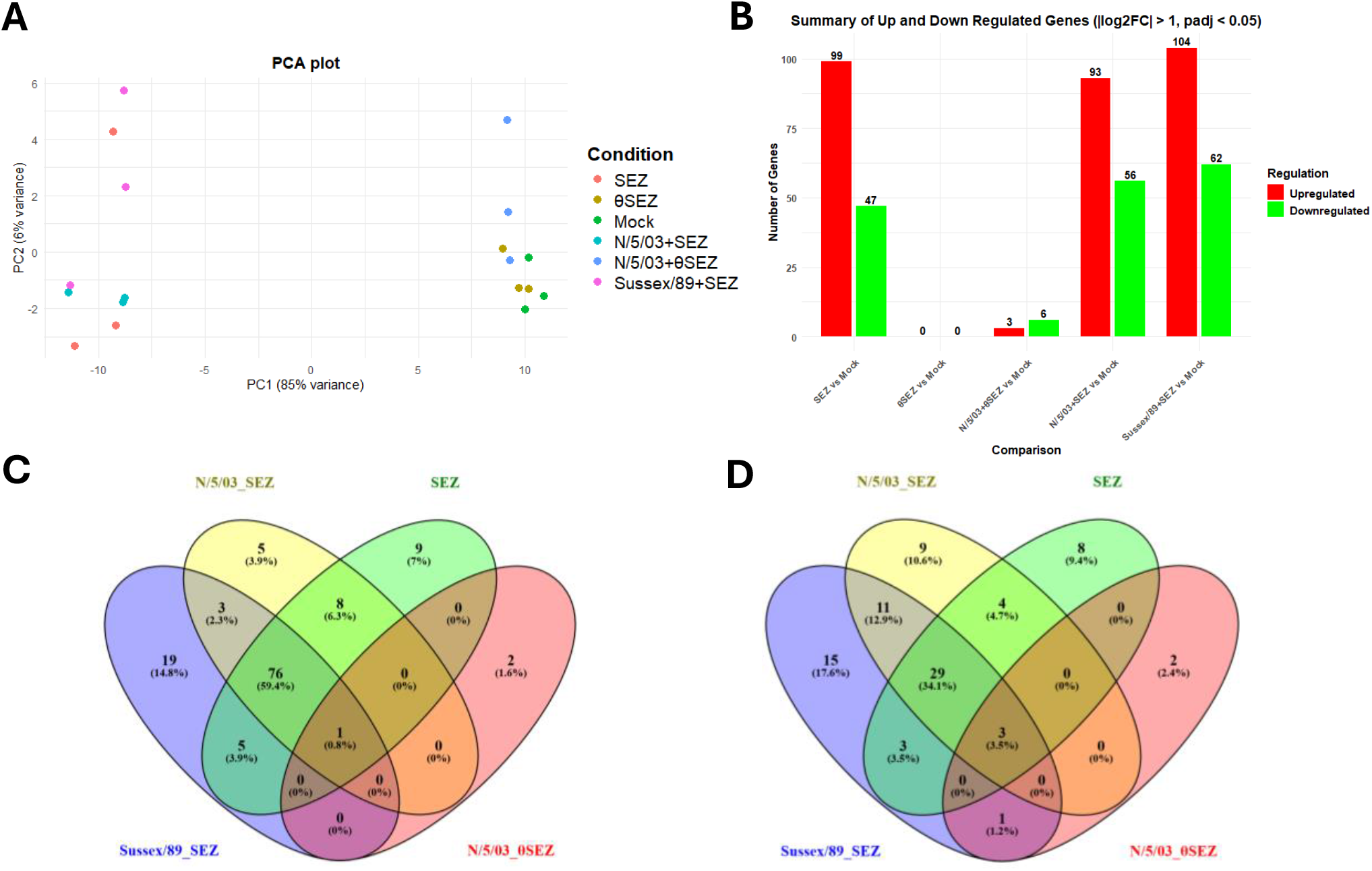
Transcriptomic analysis reveals bacterial activity-dependent responses during equine influenza A virus and *Streptococcus equi* subspecies *zooepidemicus* co-infection in DH82α cells. (A) Principal component analysis (PCA) plot showing the distribution of biological replicates (n=3 per condition) across six experimental conditions: Mock infection (Mock, green), SEZ alone (SEZ, red), heat-inactivated SEZ alone (θSEZ, yellow), N/5/03 co-infected with SEZ (N/5/03+SEZ, cyan), N/5/03 co-infected with θSEZ (N/5/03+θSEZ, blue), and Sussex/89 co-infected with SEZ (Sussex/89+SEZ, magenta). Each point represents an individual biological replicate. PC1 and PC2 explain 85% and 6% of variance, respectively. (B) Summary of differentially expressed genes across infection conditions. Bar chart showing the number of significantly upregulated (red) and downregulated (green) genes for each comparison relative to mock-infected controls. Differential expression was determined using DESeq2 with thresholds of |log_2_ fold change| > 1 and adjusted p < 0.05. (C) Venn diagram of upregulated genes showing shared and unique differentially expressed genes among N/5/03 co-infected with SEZ (N/5/03+SEZ, yellow), Sussex/89 co-infected with SEZ (Sussex/89+SEZ, blue), SEZ alone (SEZ, green), and N/5/03 co-infected with θSEZ (N/5/03+θSEZ, pink). Numbers indicate gene counts and percentages of total upregulated genes. (D) Venn diagram of downregulated genes using the same colour scheme and experimental conditions as panel C.

### 3.5 RNA-seq analysis reveals bacterial-driven transcriptional responses

To further characterise the molecular basis of transcriptional activation, detailed analysis focused on the SEZ infection condition, as this represented the core bacterial-driven response consistently observed across all infection settings. Heatmap analysis of the top 20 upregulated and top 20 downregulated genes revealed strong and reproducible clustering across biological replicates for infection with SEZ (Figure 7A). Comparative analysis revealed that the majority of these 20 differentially expressed genes were shared among SEZ alone, N/5/03 + SEZ, and Sussex/89 + SEZ infections. Specifically, 16 of the top 20 upregulated genes and 9 of the top 20 downregulated genes were common across all three conditions, confirming the presence of a conserved inflammatory transcriptional core largely driven by the bacterial component of infection. Shared upregulated genes included key pro-inflammatory and interferon-associated mediators such as IL6, IL1A, CXCL8, CXCL10, CCL3, CCL4, CCL5, CCL20, CSF2, IFNB1, PTGS2, LIF, PDCD1, PI3, and ZNF518B. In contrast, shared downregulated genes included SELENOP, MSRB1, MT1E, CNR2, ADAMTS10, CHST4, KCNIP2, GJB2, ASPM, and CYP1B1, several of which are associated with metabolic regulation, oxidative stress responses, and cellular homeostasis.

**Figure 7.**
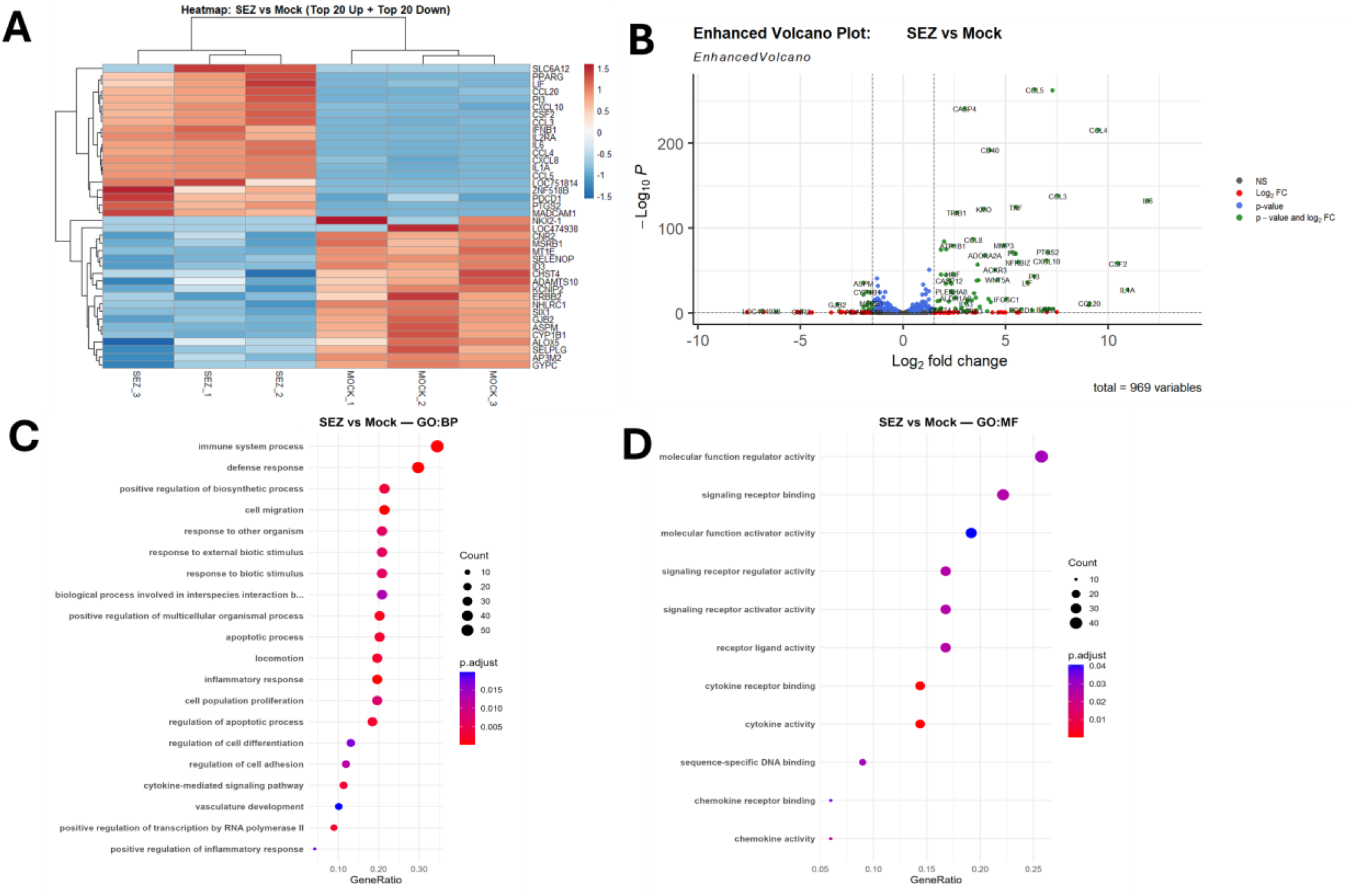
Gene expression patterns and functional enrichment analysis reveal immune activation following *Streptococcus equi* subspecies *zooepidemicus* infection in DH82α cells. (A) Heatmap showing the top 20 upregulated and top 20 downregulated genes with the highest absolute log_2_ fold changes in DH82α cells following SEZ infection compared with mock-infected controls. Rows represent individual genes; columns represent biological replicates (n=3 per condition). Colour scale indicates normalized expression levels (blue = low expression, red = high expression). (B) Enhanced volcano plot displaying differential gene expression for SEZ infection versus mock controls. The x-axis shows log_2_ fold change and y-axis shows -log_10_ adjusted p-value for all detected transcripts (total = 969 variables). Significantly differentially expressed genes meeting thresholds (|log_2_ fold change| > 1 and adjusted p < 0.05) are highlighted: upregulated genes (red), downregulated genes (blue), and non-significant genes (grey). Top upregulated genes are labelled. (C) Gene Ontology Biological Process (GO:BP) enrichment analysis of upregulated genes following SEZ infection. Dot plot shows the top significantly enriched biological process terms (adjusted p < 0.05). The x-axis represents gene ratio (proportion of genes in each term), dot size reflects the number of genes, and colour intensity corresponds to adjusted p-value significance. (D) Gene Ontology Molecular Function (GO:MF) enrichment analysis of upregulated genes following SEZ infection. Dot plot format identical to panel C, showing enriched molecular function categories.

Despite this strong overlap, subtle strain-specific differences were observed. N/5/03 and SEZ co-infection uniquely upregulated TNF and F3, whereas Sussex/89 and SEZ uniquely induced CASP8, NFKBIZ, and PSEN2, indicating additional modulation of apoptotic and NF-κB–associated regulatory pathways. Sussex/89 and SEZ co-infection also exhibited broader downregulation of regulatory genes, including HMOX1 and HPGDS. Although the core inflammatory programme was conserved across conditions, Sussex/89 and SEZ co-infection generally demonstrated a greater magnitude of transcriptional activation compared with N/5/03 and SEZ, indicating enhanced immune modulation capacity.

In contrast, heat-inactivated SEZ (θSEZ) infection did not produce detectable transcriptional changes, while co-infection with N/5/03 and heat-inactivated SEZ (N5 + θSEZ) resulted in only a small number of differentially expressed genes. These genes were not included in the heatmap, as the analysis focused on the top 20 significantly altered genes across active infection conditions.

Volcano plot analysis confirmed a clear rightward skew in gene expression changes, indicating that the majority of differentially expressed genes with higher levels of gene expression were upregulated compared to mock-infected controls (Figure 7B), consistent with the predominance of upregulated transcripts observed in the Venn diagram analysis (Figure 6C and D) and demonstrating that bacterial infection primarily activates rather than suppresses the host transcriptional response. In contrast, most downregulated genes showed more modest changes in gene expression levels. Gene Ontology enrichment analysis of upregulated genes revealed strong enrichment of immune and inflammation-related categories (Figure 7C and D), with Biological Process terms including immune system process, defence response, cell migration, inflammatory response, and cytokine-mediated signalling pathway, while Molecular Function enrichment identified cytokine activity, cytokine receptor binding, and signalling receptor binding as dominant terms. These enrichment patterns highlight the central role of cytokine signalling and immune activation in the host response to SEZ infection, with the prominent enrichment of cytokine-related functions correlating with the upregulation of multiple chemokines (CCL3, CCL4, CCL5, CXCL8) and interleukins (IL1A, IL6, IFNB1) observed in the core gene set, indicating co-ordinated activation of pro-inflammatory signalling response.

### 3.6 RT-qPCR and ELISA validation of key cytokine and interferon responses

To validate the RNA-seq findings and assess protein-level expression, RT-qPCR analysis was performed on key immune genes, followed by ELISA quantification of secreted cytokines in cell culture supernatants. RT-qPCR analysis of DH82a cells revealed upregulation of pro-inflammatory cytokines IL-6, TNF-α and IL-8 following infection with SEZ compared with mock and θSEZ (Figure 8A, B and D). Among these, IL-6 exhibited the strongest transcriptional response, approximately 1000-fold increase, while TNF-α showed a more modest increase, of around 70-fold. Importantly, no significant differences were observed between SEZ alone and SEZ following viral pre-infection with either N/5/03 or Sussex/89 for any of these pro-inflammatory cytokines.

**Figure 8.**
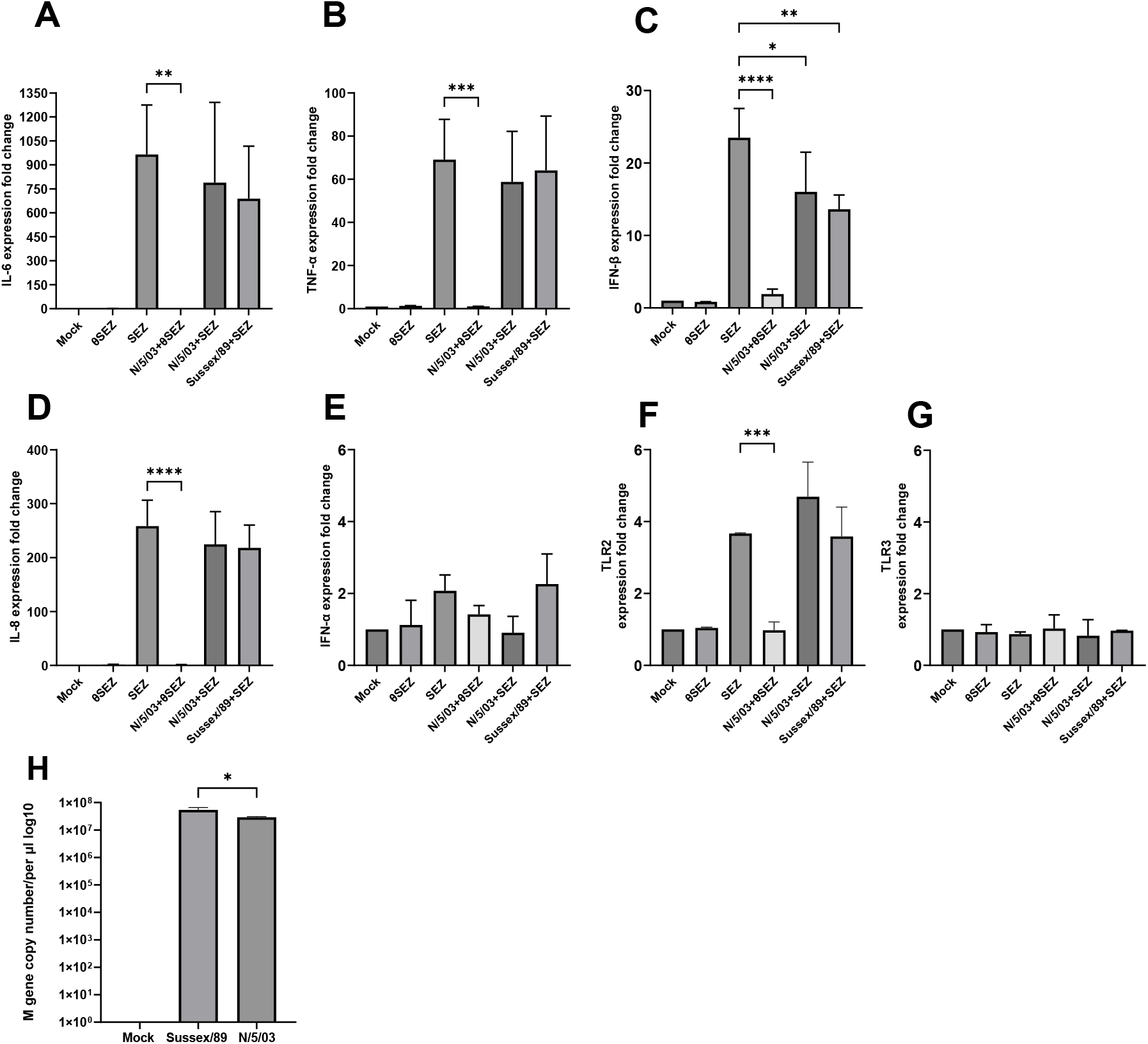
Cytokine, toll like receptor (TLR) and influenza virus matrix (M) gene expression in DH82α cells following equine influenza A and *Streptococcus equi* subspecies *zooepidemicus* (SEZ) co-infection or SEZ infection alone. RT-qPCR analysis of mRNA expression levels of (A) IL-6, (B) TNF-α, (C), IFN-β (D) IL-8, (E) IFN-α, (F) TLR2, and (G) TLR3 in DH82α cells. Cells were mock-infected or infected with SEZ or θSEZ alone, or pre-infected with N/5/03 or Sussex/89 for 24 h followed by infection with SEZ for 6 h (co-infection). Data are expressed as fold change relative to mock-infected controls (mean ± SD, n = 3). Significant differences are indicated: *p < 0.05, **p < 0.01, ***p < 0.001, ****p < 0.0001. (H) Viral M gene copy numbers quantified by RT-qPCR in DH82α cells during co-infection (Sussex/89+SEZ and N/5/03+SEZ) at 30 h post-viral infection.

Type I interferon responses differed, with IFN-α expression remaining low across all conditions (Figure 8E). In contrast to SEZ infection alone, IFN-β expression was significantly reduced following viral pre-infection, with approximately 7.5-fold and 9.9-fold lower expression observed in cells pre-infected with N/5/03 and Sussex/89, respectively (Figure 8C). Pattern recognition receptor analysis revealed significant TLR2 upregulation, of approximately 4-fold, in response to SEZ (Figure 8F), which was not significantly affected by virus co-infection. TLR3 expression remained unchanged across all conditions (Figure 8G). Viral M gene quantification in co-infected samples confirmed successful inoculum delivery and productive viral replication in DH82α cells, with comparable viral RNA levels detected between Sussex/89+SEZ and N/5/03+SEZ conditions (Figure 8H).

ELISA measurements correlated with the mRNA data for IL-6 and TNF-α, with high protein concentrations detected in SEZ-infected cultures irrespective of viral pre-infection (Figure 9A–B). No significant differences in IL-6 or TNF-α protein levels were observed between SEZ alone and SEZ following viral pre-infection with either N/5/03 or Sussex/89. In contrast, IFN-β protein levels remained at baseline (Figure 9C) despite marked transcriptional upregulation, suggesting post-transcriptional regulation during SEZ infection and EIV–SEZ co-infection.

**Figure 9.**
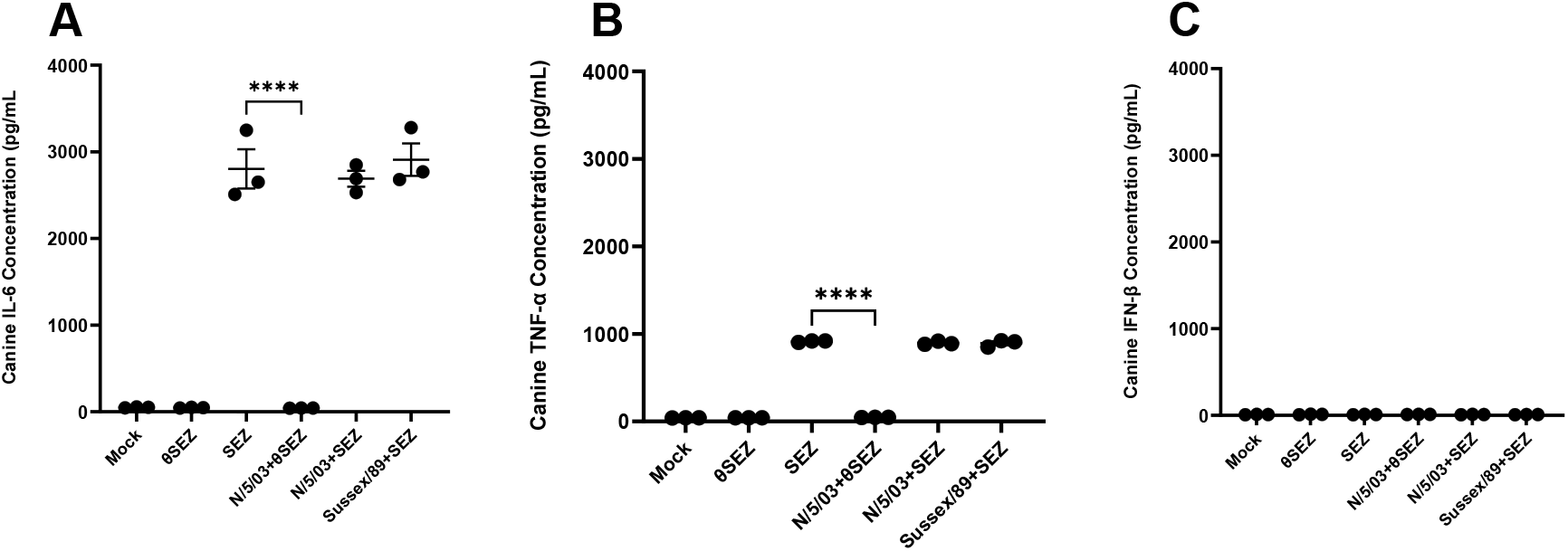
Quantification of secreted cytokine proteins in the supernatant of DH82α cells following equine influenza A and *Streptococcus equi* subspecies *zooepidemicus* (SEZ) co-infection or SEZ infection alone. Protein concentrations of (A) IL-6, (B) TNF-α, and (C) IFN-β were measured in the supernatants of DH82α cell cultures by ELISA at 30 h post-infection. Cells were mock-infected or infected with SEZ or θSEZ alone, or pre-infected with N/5/03 or Sussex/89 for 24 h followed by infection with SEZ for 6 h. Data show mean ± SEM. Significant differences are indicated: ****p < 0.0001.

## 4. Discussion

This study provides the first comprehensive characterisation of equine IAV-SEZ co-infections, revealing novel direct physical virus-bacteria interactions that fundamentally alter our understanding of the pathogenesis of secondary bacterial infection. TEM demonstrated direct binding between each of the equine IAV isolates (Miami/63, Sussex/89 and N/5/03,) and both SEZ and *S. suis*, regardless of bacterial activity, challenging prevailing models that emphasise indirect mechanisms mediated through host immune suppression [5, 45, 46]. TEM analysis revealed preferential binding of equine influenza virions to SEZ compared with *S. suis*. Although these interactions were assessed qualitatively, virion attachment to SEZ was consistently observed across multiple fields of view, while *S. suis* binding occurred less frequently.

The Miami/63 isolate was previously shown to be predominantly spherical, which is characteristic of laboratory-adapted viruses, while Sussex/89 and N/5/03 were shown to be predominantly filamentous [44]. Morphology-dependent binding patterns were observed: extensive surface contact for the filamentous isolates Sussex/89 and N/5/03 versus point-contact for the spherical virions of Miami/63.

Our lectin staining analysis revealed that SEZ shows surface expression of predominantly α2,3-linked sialic acids, while *S. suis* displays abundant α2,6-linked forms. This represents to our knowledge, the first characterisation of sialic acid linkage specificity on SEZ surfaces, while the predominance of α2,6-linked sialic acids on *S. suis* has been reported previously [9]. However, the similar binding of the equine IAV isolates to both species of bacteria suggests that the sialic acid linkage does not influence this interaction. The persistence of viral binding following neuraminidase treatment also supports the assertion that sialic acid-haemagglutinin interactions are not essential for the observed virus-bacteria association. These results suggest that equine influenza A virus-streptococcal interactions may involve more non-specific electrostatic and hydrophobic interactions mediated by viral surface proteins, ensuring stable virus-bacteria associations across diverse bacterial phenotypes with varying surface characteristics. This concept of multiple binding pathways is further supported by David et al. [6], who demonstrated high-degree binding between influenza A virus and *S. pneumoniae* even in the complete absence of pneumococcal capsular polysaccharide. The possibility of sub-optimal removal of the sialic acid components cannot be excluded; despite the use of alternate sources of sialidase/neuraminidase and repeat experiments, we could not convincingly demonstrate complete removal of the sialic acids from the bacterial cells. The difference in sialic acid expression on SEZ compared with *S. suis* raises the possibility that the bacteria have evolved specific sialic acid expression as a means of evading host recognition; this also correlates with the binding preferences for influenza virus in each species with equine viruses binding to α2,3-linked sialic acids on host cells, while swine IAVs preferentially bind α2,6-linked sialic acids, which is the predominant form expressed in the upper swine respiratory tract.

Functional studies revealed cell-type-dependent enhancement of bacterial adherence, with a significant 2-fold increase in SEZ attachment observed in DH82α cells but no enhancement in ExtEqFL cells, indicating that virus-bacteria interactions are dependent on host cell characteristics and potentially explaining anatomical differences in co-infection susceptibility within respiratory tissues. Consistent with findings by David et al. [6], who demonstrated that co-incubation of *Streptococcus pneumoniae* with human IAV enhanced viral uptake by macrophages, our SEM analysis revealed that prior equine IAV infection significantly enhanced SEZ adherence to DH82α cells. This reciprocal enhancement, where bacteria promote viral infection and viruses increase bacterial adherence, supports the existence of bidirectional synergistic interactions between respiratory viruses and bacteria. This specificity likely reflects differences in cell surface receptor expression, membrane composition, or cytoskeletal organisation that influence bacterial adherence mechanisms following viral infection [10, 11]. The differential bacterial adherence patterns observed between these cell lines may be partly explained by their distinct sialic acid receptor expression profiles, with DH82α cells exhibiting a more balanced distribution of both α2,3 and α2,6 sialic acid receptors compared to the marginal α2,6 preference observed in ExtEqFL cells (data not shown). These receptor distribution differences could influence both viral binding capacity and subsequent bacterial adherence patterns, as sialic acids serve as key attachment sites for both pathogens [47]. The increased bacterial attachment observed in DH82α cells may reflect either enhanced phagocytic clearance, as IAV-infected macrophages may demonstrate greater phagocytic activity [48, 49], or paradoxical bacterial benefit, given that influenza compromises macrophage bactericidal function through alveolar macrophage depletion and IFN-γ-mediated impairment of antibacterial responses [50-53].

Previous work using DH82α cells has demonstrated their suitability for equine influenza virus studies. Their enhanced capacity for viral replication further supports the superior experimental performance of DH82α cells. DH82α cells supported significantly higher viral titres of N/5/03 (10^5.8^ TCID_50_/ml) compared with equine lung fibroblasts (ExtEqFL; 10^4.5^ TCID_50_/ml), with ExtEqFL cells requiring higher initial viral inocula to achieve comparable levels of infection [54]. DH82α cells, derived from canine macrophages, may express surface molecules or mediate cellular signalling pathways particularly responsive to viral pre-infection, involving virus-induced changes in integrin expression, glycocalyx modification, or cytokine-mediated alterations in cell surface properties. The consistent enhancement observed for both Sussex/89 and N/5/03 suggests this effect is mediated by conserved viral components rather than strain-specific factors, supporting generalisability across equine influenza virus variants and aligning with extensive evidence from animal models demonstrating that influenza infection substantially increases bacterial loads in the respiratory tract by 25-fold or more compared to naïve hosts [5, 50].

Our transcriptomic analysis provides evidence that bacterial infection drove most transcriptional changes during co-infection with little difference in the number of differentially expressed genes (DEGs) between infection with SEZ alone (146 DEGS) or after infection with either Sussex/89 (149 DEGS) or N/5/03 (166 DEGS). The substantial gene overlap (59.4% shared across active conditions) represents a conserved bacterial response that persists during co-infection, indicating viral pre-infection does not fundamentally alter pathogen recognition or initial processing [55, 56]. This finding challenges models emphasising viral-mediated immune suppression as the primary mechanism facilitating secondary bacterial infection and supports the emerging concept that bacterial pathogens predominantly drive innate immune activation, while viral components enhance or sustain inflammatory signalling [24, 57]. These findings align with previous viral-bacterial co-infection studies showing near-identical pathway enrichment, including defence response to virus, type I interferon signalling, and innate immune response activation [23, 58], validating our transcriptomic observations across diverse cell types and model systems. However, the use of single cell cultures is unlikely to capture the complexity of intercellular interactions that occur during infection *in vivo*.

RT-qPCR analysis validated the RNA-seq findings and confirmed that SEZ is the dominant driver of pro-inflammatory activation, inducing marked upregulation of IL-6, IL-8, and TNF-α. IL-6 exhibited the strongest transcriptional response, consistent with its central role in coordinating inflammatory cascades during bacterial infections [59]. Comparable cytokine expression levels were observed for SEZ alone and following co-infection with either Sussex/89 or N/5/03, demonstrating that viral pre-infection did not amplify SEZ-induced inflammatory responses. This supports emerging evidence that bacterial pathogens dominate inflammatory signalling during viral–bacterial co-infection [21, 22, 60]. Data from RT-qPCR analysis showed no significant regulation of IFN-α or TLR3 expression under any condition and they did not appear as differentially expressed in the RNA-seq dataset. In contrast, IFN-β expression showed significantly reduced expression under co-infection conditions (7.5-fold and 9.9-fold decreases with N/5/03 and Sussex/89, respectively) in comparison with SEZ infection alone. ELISA validation confirmed elevated TNF-α and IL-6 protein in SEZ infections, consistent with transcriptional data and the dual role of IL-6 in balancing pathogen clearance with inflammation [61]. However, IFN-β displayed mRNA-protein discordance, with elevated mRNA expression but no increase in protein levels, suggesting post-transcriptional regulation or bacterial suppression of IFN-β production [62, 63].

The demonstration of direct virus-bacteria binding through multiple mechanisms suggests that co-infection may be more mechanistically complex than previously recognised, potentially requiring modified therapeutic approaches targeting virus-bacteria complexes rather than individual pathogens. The robust inflammatory responses documented, particularly dramatic cytokine upregulation driven primarily by SEZ infection, suggest that anti-inflammatory therapy may be beneficial in managing severe co-infection cases, though antimicrobial therapy remains essential given the critical role of SEZ in driving responses [60, 61]. The cell-type specificity observed in bacterial adherence enhancement indicates that therapeutic targeting may need to consider anatomical location and specific cell populations involved in infection, as virus-bacteria interactions vary significantly between different tissue types within the respiratory tract [64].

This study demonstrates that equine IAV-*Streptococcus* co-infections involve direct physical interactions mediated by multiple binding mechanisms that enhance bacterial adherence through cell-type-specific pathways, with SEZ driving robust immune activation while viral isolates provide modulatory effects. These findings represent a paradigm shift from immune suppression models to direct pathogen-pathogen facilitation strategies, establishing a molecular framework for understanding secondary bacterial complications and providing foundations for improved therapeutic and preventive strategies in equine respiratory disease.

## Author Contributions

Conceptualisation, formal analysis, review and editing: AMB, JMD and SPD. Review and editing: MAA and MM. Writing, original draft preparation, investigation, formal analysis: AKA. All authors have read and agreed to the published version of this manuscript.

## Data Availability Statement

Data are deposited in the NCBI Sequence Read Archive under BioProject accession PRJNA1454118 (SRR38177454–SRR38177459).

## Acknowledgments

The authors thank the Nanoscale and Microscale Research Centre (nmRC) at the University of Nottingham for providing access to instrumentation. The authors also thank Ms Denise McLean and Ms Nicola J. Weston for their assistance with the JEOL 2100+ TEM and the JEOL IT-200 SEM. The authors would like to acknowledge the Deanship of Graduate Studies and Scientific Research, Taif University for funding this work.

## Abbreviations

The following abbreviations are used in this manuscript:

18S: 18S ribosomal RNA
CFU: colony-forming unit
DH82α: canine macrophage-like cell line
DMEM: Dulbecco’s Modified Eagle’s Medium
ExtEqFL: equine lung cell line
FCS: fetal calf serum
HMBS: hydroxymethylbilane synthase
IAV: influenza A virus
IFN: interferon
IL: interleukin
M gene: matrix gene of influenza A
Miami/63: A/equine/Miami/1963
N/5/03: A/equine/Newmarket/5/2003
NS1: non-structural protein 1
PB1-F2: polymerase basic 1-F2
SEM: scanning electron microscopy
SEZ: *Streptococcus equi* subspecies *zooepidemicus*
Sussex/89: A/equine/Sussex/1989
TCID_50_: 50% Tissue Culture Infectious Dose
TEM: transmission electron microscopy
TLR: toll like receptor
TNF: tumour necrosis factor

